# HiCDOC: chromatin compartment prediction and differential analysis from Hi-C data with replicates

**DOI:** 10.1101/2025.09.18.677058

**Authors:** Elise Maigné, Cyril Kurylo, Matthias Zytnicki, Sylvain Foissac

## Abstract

**Motivation:** The spatial organization of the genome plays an essential role in regulating cellular functions, with A/B chromatin compartments reflecting broad differences in transcriptional and epigenetic activity. Hi-C enables genome-wide identification of such compartments, but robust differential analysis between groups of samples remains challenging. Existing approaches largely rely on Principal Component Analysis, which, applied on Hi-C matrices separately, requires heuristic sign choices to merge results and does not naturally incorporate replicates.

**Results:** Here we present HiCDOC, a Bioconductor package for the prediction and differential analysis of chromatin compartments from Hi-C data with replicates. HiCDOC uses constrained *k*-means clustering to jointly analyze multiple Hi-C matrices, incorporating replicate information to enhance robustness, and provides empirical statistical support for predicted compartment switches.

Applied to Hi-C datasets from human tissues and mouse cell lines, HiCDOC identified biologically relevant compartment changes supported by transcriptional differences. Comparisons with existing tools showed both overlap and complementarity, while a controlled benchmark with artificially introduced changes confirmed high sensitivity. Although extensively tested on pairwise comparisons, HiCDOC offers a flexible framework compatible with more complex designs and, in principle, with more than two compartment states.

By combining replicate-aware clustering, automatic A/B assignment across chromosomes, extensive quality control, and statistical evaluation, HiCDOC provides an alternative and complementary approach to PCA-based methods for compartment analysis. HiCDOC thus expands the methodological toolkit for exploring 3D genome dynamics and its role in cellular processes.

**Availability:** HiCDOC is implemented in R and C++, and is available on Bioconductor: https://bioconductor.org/packages/release/bioc/html/HiCDOC.html

## Introduction

The three-dimensional (3D) organization of the genome in the nucleus plays a crucial role in regulating essential functions of eukaryotic cells (Oudelaar and Higgs, 2020; Spielmann et al., 2018). It has been shown to be involved in cancers (Johnstone et al., 2020; Rhie et al., 2019; Xu et al., 2022), cell differentiation (Bonev et al., 2017; Oudelaar et al., 2020), and development (Lupiáñez et al., 2015; Marti-Marimon et al., 2021). The 3D conformation of chromosomes is organized by specific structural features at different scales, including DNA loops, Topologically Associating Domains (TADs), and chromatin compartments (Rao et al., 2014; Rowley and Corces, 2018). These compartments are characterized by distinct chromatin states that exhibit different epigenetic properties and gene expression patterns, typically classified into two types. Generally, “A” compartments are associated with more active histone modification marks, chromatin accessibility, and expressed genes compared to “B” compartments (Lieberman-Aiden et al., 2009). As epigenetic components of the genome, chromatin compartments are dynamic, implying that the compartment type of a given genomic position can change over time. Consequently, comparing chromatin compartmentalization between different biological groups of cells can reveal valuable information on the functional role of 3D variations of the genome (Bonev and Cavalli, 2016).

Identifying chromatin compartments, *i.e*., determining the compartment type of each genomic position, can be achieved by analyzing data from genome-wide Chromatin Conformation Capture (Hi-C) experiments (Lieberman-Aiden et al., 2009). From a given biological sample, the Hi-C experiment produces pairs of sequencing reads from interacting genomic regions. A standard analysis pipeline processes these reads to generate an interaction matrix, which shows the frequency of observed interactions between pairs of genomic regions, or “bins.” The bin size of a matrix defines its resolution. Due to the proximity-dependent ligation principle of Hi-C, interaction counts serve as a proxy for the spatial proximity of associated genomic regions. The Hi-C matrix often displays a strong signal along the diagonal and a plaid pattern, similar to a chessboard with variably sized squares. This pattern is a characteristic signature of chromatin compartments.

The traditional method for detecting this signal involves a Principal Component Analysis (PCA) of the distance-normalized (“observed/expected”) interaction matrix. Based on dimensionality reduction, this approach usually classifies genomic loci into A and B compartments based on the sign of the first eigenvector (PC1) values (Lieberman-Aiden et al., 2009). Despite their popularity, PCA-based methods have several limitations:

- As they perform dimensionality reduction, PCA-based approaches rely on the strong assumption that all relevant information can be retrieved in a single eigenvector of the entire interaction matrix.
- PC1 may not always be the most suitable eigenvector for identifying compartments, depending on factors such as the chromosome, species, or data resolution (Lieberman-Aiden et al., 2009). This often requires a tedious, case-by-case inspection of the results for each chromosome and dataset to confirm that PC1 is the most appropriate choice (Kai et al., 2023; Kalluchi et al., 2023; Rahman et al., 2023).
- The signs of the eigenvectors are arbitrarily assigned by a distinct PCA for each matrix. This makes it impossible to directly and consistently assign “A” or “B” compartment types to either the positive or negative PC1 sign (Kalluchi et al., 2023). To achieve consistent A/B labels across different chromosomes and datasets, it is necessary to “synchronize” the assignment using external genome annotation data, such as gene density, GC content, or epigenetic marks (Chakraborty et al., 2022; Lieberman-Aiden et al., 2009). In practice, this information may not be always available in the required format for the species or tissue of interest. Additionally, depending on the computational tool, this synchronization process may require a manual, per-chromosome check to decide if compartment types need to be swapped.
- Despite these known limitations and the lack of a reliable, automated method for consistent compartment type assignment, most available PCA-based tools do not include dedicated quality control (QC) procedures to assess the validity of these choices and processes.

Comparative analysis of chromatin compartmentalization is even more challenging. Most comparative studies to date use a two-step qualitative approach: they first predict compartments for each biological group, often by merging replicates into a single matrix, and then compare the predictions. While straightforward and widely used, this approach has no statistical support and is heavily influenced by factors such as matrix resolution, dataset size, and intra-or inter-group variability (Kalluchi et al., 2023; Marti-Marimon et al., 2021). Some studies have adopted more quantitative approaches, comparing compartment-related metrics often derived from PCA (Dixon et al., 2015; Narang et al., 2023; Rahman et al., 2023). However, these approaches are typically implemented as “in-house” scripts designed for a specific project, and do not provide a generic computation tool for broader a usage and evaluation.

Nonetheless, a few statistical methods have been specifically developed and implemented to identify significant differences in compartmentalization between groups of matrices. HOMER (Heinz et al., 2018) performs a PCA and compares PC1 values from different conditions using a gene expression analysis based on the limma R package (Ritchie et al., 2015). The choice of the PC and of its sign has to be manually checked and potentially corrected for each individual matrix. Additionally, HOMER is not compatible with standard Hi-C matrix formats and requires the completion of its entire Hi-C workflow beforehand, making it challenging to analyze existing matrices. DcHiC (Chakraborty et al., 2022) also adopts a PCA approach, where PC values between conditions are quantile-normalized and compared using the Mahalanobis Distance to detect atypical bins and assess statistical significance. Unlike HOMER, dcHiC automatically determines which PC to use and how to orient it based on GC content and gene density, requiring users to provide this information. DARIC (Kai et al., 2023) is the only differential method that does not rely on PCA. Instead, it calculates a Preferential Interaction Score (PIS) for A vs. B compartment at each bin and compares them between conditions to identify differences using a Hidden Markov Model. Statistical significance is determined by comparing replicates from the same group to estimate an expected background. However, DARIC requires a pre-existing compartment annotation to operate, making it dependent on other predictors to assign compartment types. Moreover, its current version can only be used with human and mouse genomes (mm9, mm10, hg19, hg38), which significantly limits its applicability to the broader animal genome research community (Cheng et al., 2024; MacPhillamy et al., 2021).

In order to address these shortcomings and to provide an alternative method, we present HiCDOC, a Bioconductor package dedicated to the differential analysis of chromatin compartmentalization from Hi-C data. HiCDOC features several unique properties. Unlike PCA-based methods that rely on comparing the results of independent dimensionality reductions, HiCDOC performs a *k*-means clustering of the bins from all matrices for each chromosome, leveraging the consistency between replicates. In addition, HiCDOC uses interaction values to automatically assign A and B compartment types for all bins and matrices, eliminating the need for external annotation data. HiCDOC also uses several QC metrics to automatically detect potential artifacts, and an empirical *p*-value estimation method to assess the statistical significance of the results. Last, HiCDOC is implemented as an R Bioconductor package, enabling easy integration with existing workflows and data structures.

We demonstrate the effectiveness of HiCDOC in identifying significant compartment differences between various cell types and experimental conditions. Comparisons with existing tools highlight both the relevance and the complementarity of HiCDOC’s results on public datasets. By implementing an alternative method for chromatin compartment analysis, HiCDOC provides a new approach to investigate the intricate spatial organization of the genome and its role in cellular processes.

## Materials and Methods

### Differential compartment analysis with HiCDOC

#### Method overview

The aim of HiCDOC is to identify significant differences of chromatin compartmentalization along the genome between groups of Hi-C matrices. Each group typically corresponds to a biological factor of interest (or “condition”), and contains one matrix per replicate. Most studies compare two groups. Broadly, HiCDOC first assigns a compartment type A or B to each genomic bin for each group, assuming that the compartmentalization status should be the same for all replicates of the same group. Bins with a different compartment type between groups (A vs. B) are considered as candidate “switches.” To identify true switches, a statistical significance is computed and assigned to each candidate using an empirical background, assuming that most of the genome does not change compartment between conditions. The processing steps are the following.

1. Data loading and preprocessing
2. Normalization
3. Compartment prediction
4. Quality Control and visualization

#### Data loading and preprocessing

Briefly, the package first reads a list of matrices encoded in.hic,.cool,.mcool, or tabular HiC-Pro file format, and a metadata file describing the experimental design, *i.e*., which biological group is assigned to each matrix. The first step performs data cleaning and filtering. It removes short chromosomes (less than 100 bins by default), sparse replicates (less than 30% positive interaction counts), and bins with few interactions (less than one interaction on average). The parameter settings can be changed if necessary.

#### Data normalization

Data normalization can be decomposed into three steps. First, an inter-matrix normalization is performed in order to account for technical biases like sequencing depth discrepancies across libraries. For this, HiCDOC applies a cyclic loess normalization, implemented in Stansfield et al. (2019), in order to jointly normalize matrices from various groups and replicates. Then, to account for biases like repeat or restriction site density variations along the sequence, an intra-matrix normalization is performed by applying the KR algorithm to each matrix (Knight and Ruiz, 2012), so that all the bins end up with the same total number of interactions. Last, an observed/expected MD-loess (Stansfield et al., 2018) normalization allows to obtain comparable interaction values regardless of the distance to the diagonal.

#### Compartment prediction by constrained clustering

To predict the compartment type of each bin in each matrix, HiCDOC performs a constrained *k*-means clustering (Wagstaff et al., 2001) of the vectors of interaction values, processing each chromosome separately. The clustering considers two clusters (*k* = 2) per biological condition to represent the A and B compartments, iteratively assigning each vector to the closest centroid. As a specific property of the constrained *k*-means clustering, vectors from replicates of the same condition have to be assigned to the same centroid. The assumption behind this constraint is that intragroup variability should not exceed intergroup variability. At the end of the clustering, each bin of each matrix is assigned to one of the two clusters.

Then, each compartment type (A or B) is assigned to a centroid. Instead of relying on user-provided external data, HiCDOC uses an empirical property of the matrices based on a new metric: the *Self-Interaction Ratio* (SIR). For each bin of each matrix, the SIR is the number of interactions between the bin and itself, divided by the total number of interactions in the bin. The SIR therefore represents the proportion of interactions along the diagonal of the matrix. Since the matrices are KR-normalized at this stage, the diagonal value can simply be used directly in practice. The cluster with the highest median SIR is labeled compartment A and the other one compartment B. Consequently, all matrices feature a sequence of A/B compartment predictions, defining two types of bins: stable bins, with the same compartment label in both conditions, and switching bins, with different compartments.

For each bin of each matrix, HiCDOC computes the *concordance*, a number between *−*1 and +1 that reflects the relative distance between the corresponding interaction vector and both centroids. The concordance of a bin *x*, considering centroids *c*_A_ and *c*_B_ of compartments A and B respectively, is

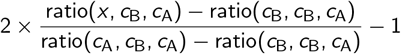

with ratio 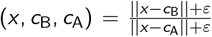, and *ε* a small value (10^*−*10^ in the implementation) to avoid divisions by zero. A concordance of *−*1 means that the bin is very similar to the centroid of compartment B, while a concordance of +1 indicates a close proximity to compartment A.

In order to obtain a background distribution under the null hypothesis (*i.e*. no compartment switch), the concordances of stable bins are considered. For each stable bin, we compute the median concordance in each group, then the difference between these medians. This distribution is then used to assess the statistical significance of the switching bins. Resulting *p*-values are subsequently adjusted for multiple testing using the Benjamini–Hochberg procedure (Benjamini and Hochberg, 1995).

#### Quality controls and visualizations

HiCDOC automatically performs three quality controls on each chromosome to assess the consistency of the clustering process.

The first step focuses on how the centroids ended up distributed in the subspace. If normalization and clustering provided expected results, centroids of the same type should be close to each other, *i.e*. the compartment type should be more discriminant than any other factor. To control for this, a PCA is performed on the centroids. The resulting first Principal Component must fulfill two conditions: (1) its sign should be the same for centroids of the same type, and (2) its inertia should account for at least 75% of the PCA dispersion. Otherwise, the chromosome gets flagged by the QC process.

The last control focuses on the relevance of the A/B label assignment and on the actual discrepancy between the two compartment types. We gathered the SIR values of the bins assigned in the A compartment, the SIR values of the bins assigned in the B compartment, and compared the two distributions with a Wilcoxon test. If no significant difference is found, the chromosome is flagged. This control helps to avoid improper segmentation of the chromosome, not driven by biological condition but by another factor such as the chromosome arm, for example (Lieberman-Aiden et al., 2009).

#### Implementation

The HiCDOC package is developed in R and is available as a Bioconductor package (Gentleman et al., 2004): https://bioconductor.org/packages/release/bioc/html/HiCDOC.html. As such, it is fully compliant with the Bioconductor’s data structures. In addition, while each step of the analysis can be executed independently, a global function allows to run the entire workflow as an integrated pipeline.

### Benchmark

#### Evaluated tools

We compared HiCDOC version 0.99.15 with HOMER version 4.11 (Heinz et al., 2018), DARIC version 0.2.20 (Kai et al., 2023), and dcHiC v2.0 (Chakraborty et al., 2022). All tools were used with their default options.

Unlike the other tools, which can process interaction matrices, HOMER needs to run an entire Hi-C data workflow, starting from raw reads. These reads are trimmed, mapped, and aggregated to build the interaction matrices. Each matrix is normalized, first by library size, then by distance from the diagonal (observed/expected). A Principal Component Analysis is then used to segment the chromosome into compartments, usually using the first eigenvector PC1. The correct sign of the PC1 to be used for A/B assignation can be inferred using user-provided annotation data, like histone marks or Transcription Start Sites.

While DARIC can process distance-normalized interaction matrices, it needs a corresponding A/B compartment prediction, obtained by another tool. For each bin, it computes a Preferential Interaction Score (PIS), which is the log2-transform of the average interaction with a bin in the A compartment, divided by the average interaction with a bin in the B compartment. This PIS is smoothed using a gaussian filter and normalized using a robust rescaling of the MA-plot. A Hidden Markov Model then segments the genome into four states: strong “A to B”, weak “A to B”, and the converse for the changes “B to A”. Last, a null distribution of the difference between PIS from replicates is used to provide a *p*-value for the strong changes found in the previous step.

dcHiC starts from an interaction matrix, which is processed through a PCA. The genomic GC content is used to assign A/B compartments. A multivariate distance measure, the Mahalanobis distance, is computed using the quantile-normalized eigenvectors from all the matrices. A chisquare test on this distance identifies significant changes, using variance across replicates as covariates for independent hypothesis weighting.

#### Evaluation data

Three datasets were used to benchmark the tools.

- The human Hi-C datasets were generated by the ENCODE consortium (The ENCODE Project Consortium, 2012) and were downloaded from the ENCODE data portal (Luo et al., 2019) at https://www.encodeproject.org/ with the following identifiers: ENCBS393MMT, ENCBS476FKF, ENCBS565LDI, ENCBS564MPZ, ENCBS836DBQ, ENCBS494DUH. These data come from Hi-C experiments that were performed on samples from two tissues: transverse colon and skeletal muscle (gastrocnemius medialis), each with 3 biological replicates (different donors). To obtain Hi-C matrices, raw sequencing reads were processed using the nf-core/hic pipeline (Ewels et al., 2020) v1.2.2 on the assembly version GRCh38 of the human genome using the following arguments: --min_mapq 10 --restriction_site ’^GATC’ --ligation_site ’GATCGATC’ --min_insert_size 20 --max_insert_size 1000 --rm_singleton --rm_dup --skip_ice --bwt2_opts_end2end --very-sensitive -L 30 --score-min L,-0.6,-0.2 --end-to-end --reorder –bwt2_opts_trimmed --very-sensitive -L 20 --score-min L,-0.6,-0.2 --end-to-end --reorder. Hi-C matrices were generated at a 200 kb resolution. Gene expression values from RNA-seq experiments were also retrieved for the same samples on the ENCODE data portal. A mean expression value was assigned to each bin per tissue by averaging the TPM values of all genes within the bin across replicates. These values were then used to compute a logFC ratio between tissues.
- The murine Hi-C datasets come from a neural development study (Bonev et al., 2017) and were downoladed from the SRA (https://www.ncbi.nlm.nih.gov/sra) using the accession ID SRP101791 (GEO: GSE96107). To obtain Hi-C matrices, raw sequencing reads were processed using the nf-core/hic pipeline (Ewels et al., 2020) v1.3.0 on the assembly version GRCm39 of the mouse genome using the following arguments: --min_mapq 10 --digestion ‘dpnii’ --min_insert_size 20 --max_inser_size 1000 --bwt2_opts_end2end --very-sensitive -L 20 --score-min L,-0.6,-0.3 --end-to-end --reorder --bwt2_opts_trimmed -5 5 --very-sensitive -L 20 --score-min L,-0.6,-0.3 --end-to-end --reorder. Samples correspond to 3 cellular differentiation stages during neuronal development: mouse embryonic stem cells (mESC), neural progenitor cells (NPC), and cortical neurons (CN), with 4 replicates per cell type. In line with the comparison made in the dcHiC article, we chose to compare ESC and NPC, at 100 kb resolution. Gene expression values from Chakraborty et al., 2022 were downloaded from GEO using accessions GSM2533843-48 and processed as described for the human data.
- The semi-simulated data was generated from the mouse data (see above), introducing artificial compartment modifications at known positions to have a ground truth. First, consistent predictions from both dcHiC and HiCDOC were considered, focusing on compartment transitions, *i.e*. bin pairs where bin *i* was assigned to one compartment type by both tools in both conditions, and bin *i* + 1 was assigned to the other compartment type by both tools in both conditions. For all replicates of the second condition, the interaction vector of bin *i* was copied into the bin *i* + 1 to artificially shift the compartment transition by one bin downstream. In addition, this new *i* + 1 vector was also shifted by one bin along the *j* axis to preserve the diagonal structure of the matrix. As a consequence, the bin *i* + 1 should not be assigned to the same compartment type in both conditions, producing a true positive target. A total of 49 compartment transitions were modified in this way.

## Results

In order to assess HiCDOC’s performances, we ran it on real publicly available data from two distinct studies: one from the ENCODE project, comparing human tissue samples from muscle and colon, and one from a neural cell differentiation study in mouse, comparing embryonic stem cells (ES) and neuronal progenitors (NP). We also processed these datasets with three state-of-the-art tools for differential compartment calling: HOMER, DARIC and dcHiC.

In addition, we performed a complementary test using real but edited data in order to compare the best predictors in the context of a controlled setting (“semi-artificial data”). Details about the datasets and the tools are provided in the Materials and Methods section.

### HiCDOC identifies consistent compartment switches in real data from human and mouse

In both human and mouse data sets and regardless of resolution, the number of predicted compartment differences between conditions varies widely between the tools (Figure 1).

**Figure 1.**
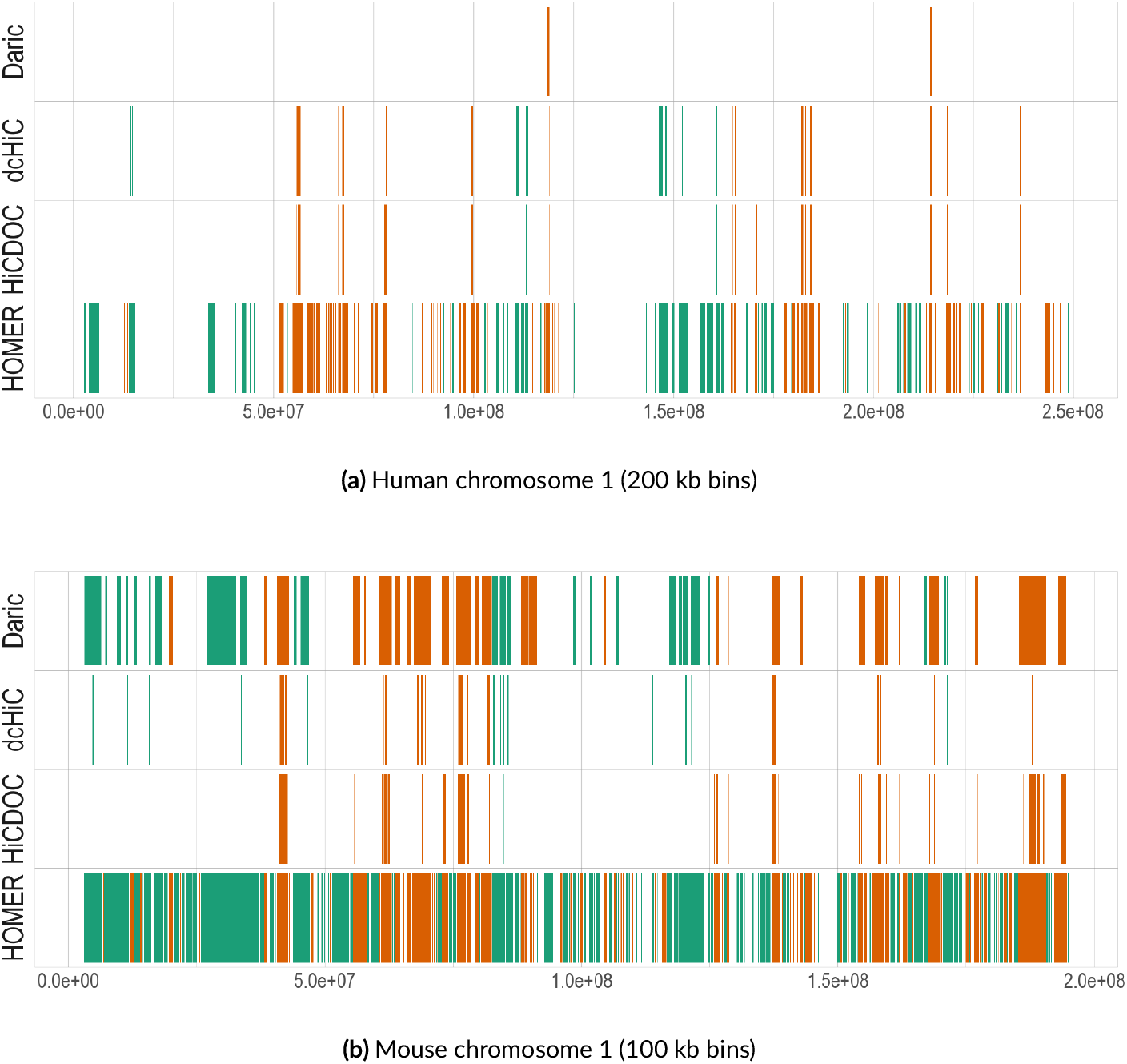
Compartment switch predictions along the first chromosome of each species between conditions: muscle vs. colon tissue samples in human (top) and ESC vs. NPC cell lines in mouse (bottom). Horizontal tracks show AB (green) or BA (orange) predicted switches according to Daric, dcHiC, HiCDOC and HOMER along the first chromosome (x-axis: genomic position in bp).

In general, HOMER produced the highest number of predicted switches for both species, with 4, 049 switching bins of 200 kb in human and 14, 593 bins of 100 kb in mouse, representing about 27% and 49% of the genome, respectively (Figure 1, 2). DARIC predicted 1, 015 human and 7, 476 mouse switches, dcHiC 1, 024 and 1, 240, and HiCDOC 765 and 1, 696 (Figure 2).

**Figure 2.**
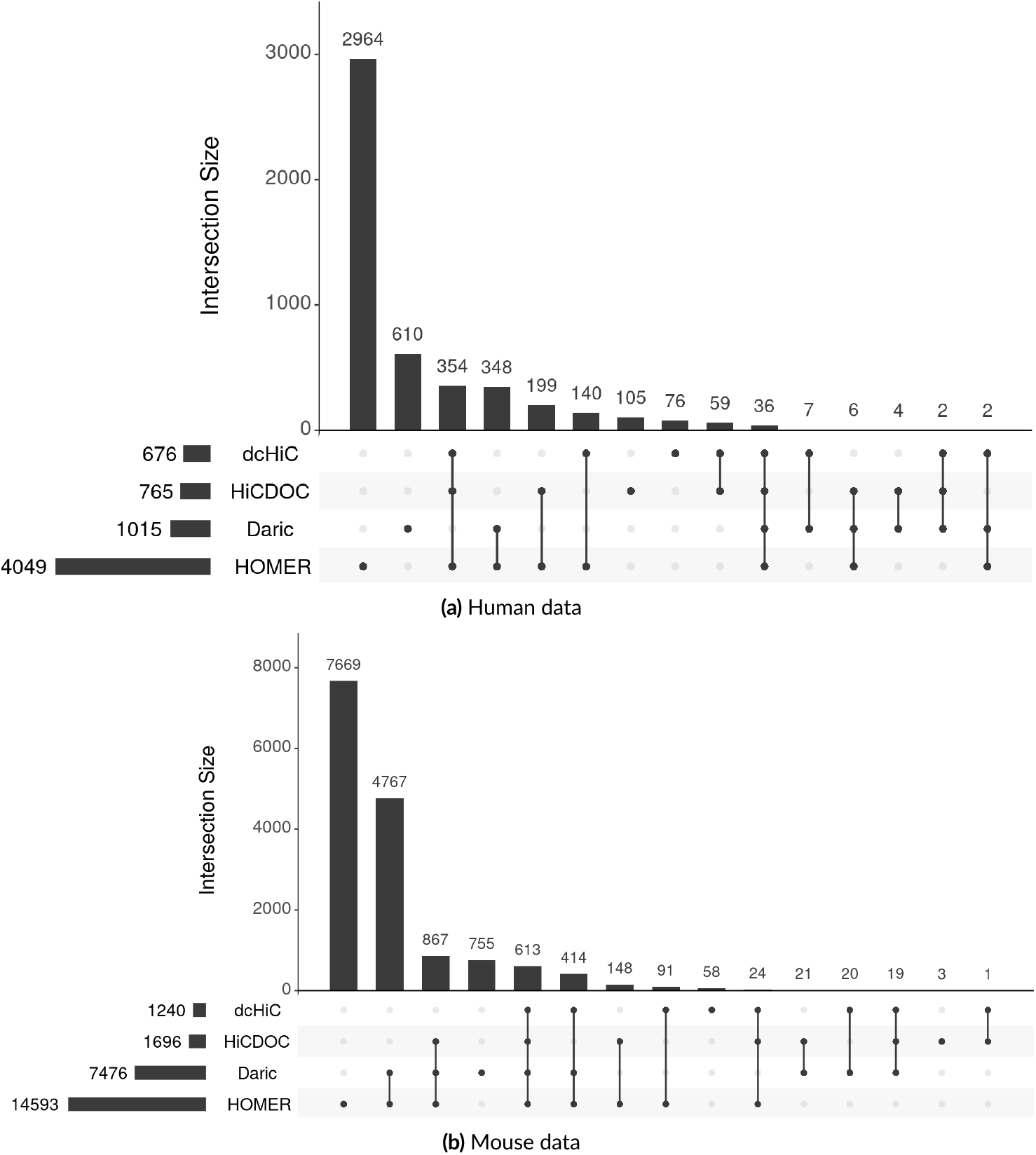
Number of significant compartment differences predicted from each tool for human (top) and mouse (bottom). Horizontal bars on the left indicate the total number of switching bins predicted by each tool, while vertical bars represent common or tool-specific predictions.

As true compartment switches are not known for these datasets, we evaluated the consistency between tools by comparing their predictions, assuming that predictions that are supported by several tools are more likely to indicate genuine compartment switches compared to tool-specific predictions. Prediction consistency also varies across tools and datasets (Figure 2). The highest proportion of tool-specific predictions was generated by HOMER, with a large majority of its predictions not supported by any of the other tools (68% in human 53% in mouse). On the other side of the spectrum, HiCDOC seems to produce the most consistent results: unlike the other tools, most of the HiCDOC predictions are supported by at least two other tools (60% in human and 90% in mouse), with dcHiC closely following (45% and 86%, respectively). This proportion is much lower for the other tools: from 6% to 25% for DARIC and from 11% to 13% for HOMER, suggesting more consistent results from HiCDOC and dcHiC. In addition, HiCDOC produced the smallest number of tool-specific predictions in both species (3 switches in mouse and 98 in human). These results support the reliability of HiCDOC’s predictions.

### HiCDOC produces biologically relevant results

Although no ground truth is available for these datasets, compartment switches are known to be associated with changes in gene expression. On average, chromatin accessibility tends to be positively correlated with gene expression in several species (Dixon et al., 2015; Foissac et al., 2019). Therefore, in order to assess the biological relevance of the predictions, we computed the average difference of gene expression as logFC values between conditions in each predicted switch, considering “A*→*B” (AB) and “B*→*A” (BA) switches separately. In principle, the resulting distributions of the logFC expression values should differ between AB and BA switches. Remarkably, this is the case for all tools, each producing significantly different distributions in human and mouse (*p*-values *<* 3.10^*−*3^, Wilcoxon test, Figure 3). However, in both species, the difference between the median logFC values is higher in the case of dcHiC and HiCDOC than in the case of HOMER and DARIC (Figure 3). Furthermore, considering the asymmetry of the distributions, we computed for each tool the proportion of predicted AB and BA switches with negative and positive logFC values, respectively. Again, this proportion is higher for HiCDOC and dcHiC than for HOMER and DARIC (Table 1), suggesting a higher consistency with expression data for the tools. Taken together, these results strongly support the biological relevance of the results from HiCDOC and dcHiC.

**Table 1.**
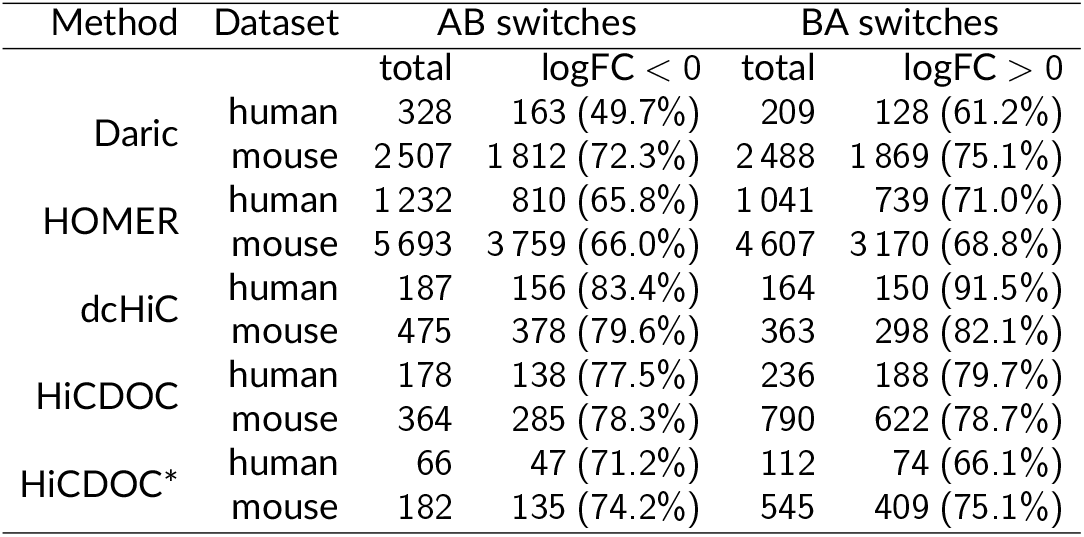
Number of 200 kb (human) and 100 kb (mouse) genomic bins with a predicted compartment switch between conditions according to each tool. Proportions of bins with a gene expression difference of the expected sign (negative and positive logFC for AB and BA switches, respectively) are also indicated. HiCDOC* is the set of predictions given by HiCDOC, but not dcHiC.

| Method | Dataset | AB switches |  | BA switches |  |
| --- | --- | --- | --- | --- | --- |
|  |  | total | logFC < 0 | total | logFC > 0 |
| Daric | human | 328 | 163 (49.7%) | 209 | 128 (61.2%) |
|  | mouse | 2 507 | 1 812 (72.3%) | 2 488 | 1 869 (75.1%) |
| HOMER | human | 1 232 | 810 (65.8%) | 1 041 | 739 (71.0%) |
|  | mouse | 5 693 | 3 759 (66.0%) | 4 607 | 3 170 (68.8%) |
| dcHiC | human | 187 | 156 (83.4%) | 164 | 150 (91.5%) |
|  | mouse | 475 | 378 (79.6%) | 363 | 298 (82.1%) |
| HiCDOC | human | 178 | 138 (77.5%) | 236 | 188 (79.7%) |
|  | mouse | 364 | 285 (78.3%) | 790 | 622 (78.7%) |
| HiCDOC* | human | 66 | 47 (71.2%) | 112 | 74 (66.1%) |
|  | mouse | 182 | 135 (74.2%) | 545 | 409 (75.1%) |

**Figure 3.**
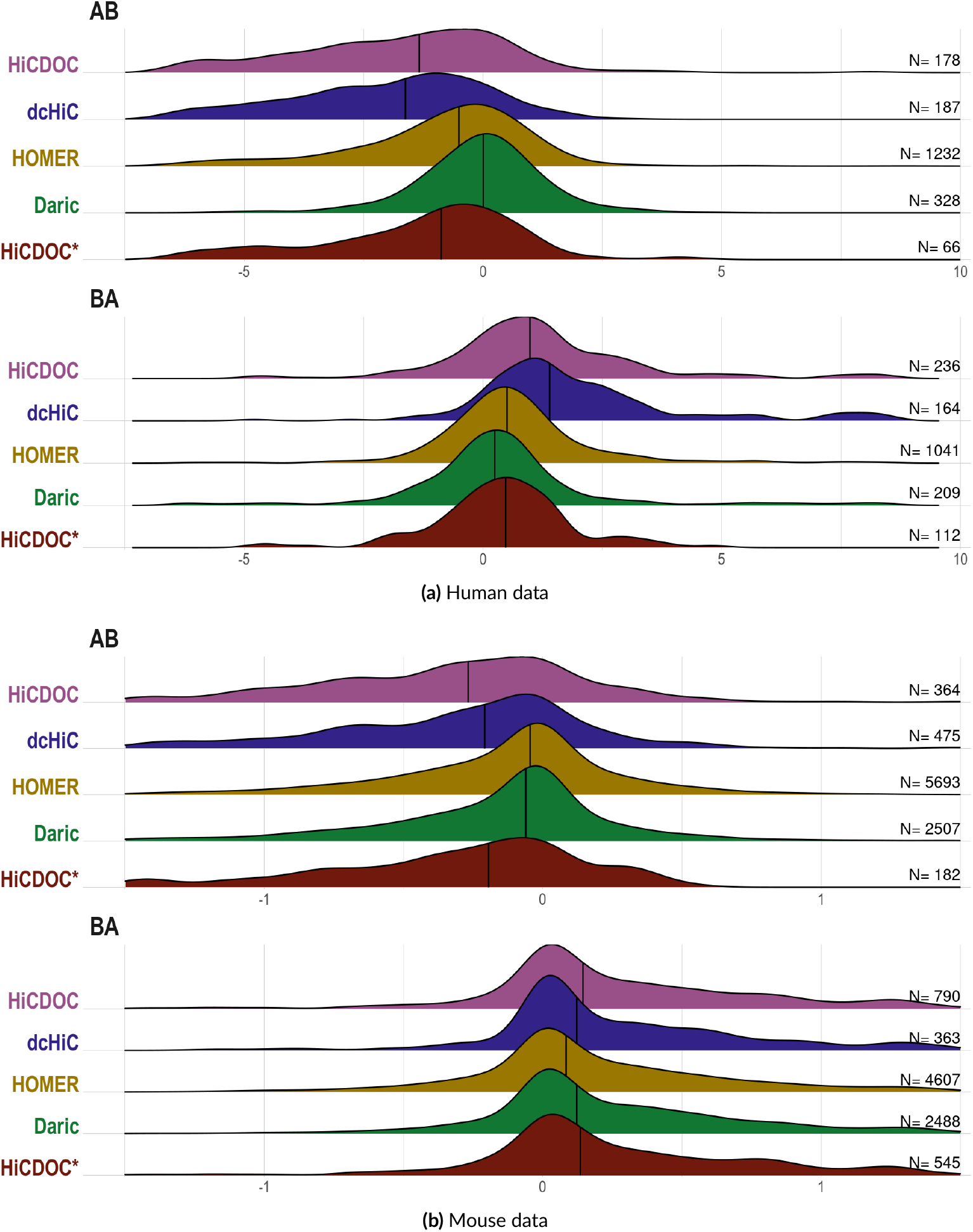
Differential gene expression in predicted compartment switches. Distributions show the average differential gene expression values (logFC, x-axis) of the genes located in bins with a predicted AB or BA compartment switch between conditions: muscle vs. colon for human (top) and ES vs. NP for mouse (bottom). Numbers of switching bins are indicated on the right, and median values by vertical bars. Genomic regions with a compartment switch from A to B are expected to show chromatin compaction and negative logFC values on average, while the opposite is expected for AB opening regions.

Finally, we sought to assess the specific contribution of HiCDOC predictions compared to those from dcHiC. For this, we focused on differential gene expression results in HiCDOC switches that were not predicted by dcHiC, ignoring the overly abundant predictions of DARIC and HOMER (Figure 3 and Table 1, HiCDOC* predictions). Similarly to the entire set of predictions, we obtained a significant difference between distributions of logFC differential expression values in AB vs. BA HiCDOC-specific switches (*p*-value *<* 10^*−*8^ in human and *<* 10^*−*38^ in mouse, Wilcoxon test). Also, HiCDOC-specific logFC distributions showed the same expected asymmetry pattern, as previously observed for the entire set, with a prevalence of positive and negative logFC expression values in BA and AB switches respectively (Figure 3 and Table 1). These results highlight the specific value of HiCDOC compared with dcHiC, emphasizing the importance of complementary analysis methods for chromatin structure studies.

### HiCDOC performs well on a dataset with ground truth

In order to perform an evaluation using a known ground truth, we edited the *Mus musculus* Hi-C matrices. In brief, all consistently predicted compartment transitions between adjacent bins were artificially shifted by one bin downstream in each replicate of one condition, generating 49 artificial switches created from real data (see Methods). We then processed this edited dataset with HiCDOC and dcHiC.

Among the 49 newly introduced compartment differences, dcHiC identified 29 of them as differential (59.2%). HiCDOC, on the other side, correctly predicted all 49 of them as differential. This results highligths HiCDOC’s capacity to accurately detect differences in chromatin compartmentalization.

## Discussion

In this report, we presented HiCDOC, a method for the identification and differential analysis of chromatin compartments from Hi-C data. By integrating replicate information into a constrained clustering framework, HiCDOC provides a robust basis for consistent compartment assignments. Unlike PCA-based methods, which process Hi-C matrices separately and rely on heuristic sign choices to merge results, HiCDOC jointly analyzes all replicates and experimental groups from the normalization step onward, accounting for inter-matrix biases and automatically synchronizing compartment types across chromosomes. In addition, by comparing candidate switches with a background distribution derived from stable bins, HiCDOC provides an empirical framework to assess statistical significance, avoiding strong assumptions about the underlying data distribution. Together, this design complements existing approaches and offers an alternative perspective on compartment organization.

Applied to Hi-C datasets from human and mouse, HiCDOC identified consistent and biologically relevant compartment switches. Notably, HiCDOC predicted fewer tool-specific differences than other state-of-the-art methods, supporting the robustness of its results. Moreover, HiCDOC predictions showed concordance with gene expression changes between conditions, reinforcing their biological relevance. Importantly, although HiCDOC and dcHiC often converged on similar results, HiCDOC-specific predictions also corresponded to expression differences, suggesting that non-PCA-based methods can provide complementary insights into chromatin dynamics. Considering the large variability in predictions across tools and datasets, these findings underline the importance of developing alternative methods and applying multiple computational approaches when analyzing chromatin structure data, as already exemplified in the context of TAD and loop detection (Dali and Blanchette, 2017; Liu et al., 2023; Zufferey et al., 2018).

Evaluation on a semi-artificial dataset with a controlled ground truth further highlighted the capacity of HiCDOC to recover experimentally introduced compartment changes with high sensitivity and precision. While this specific setting only evaluates the capacity of the methods to identify a limited number of targets, semi-artifical datasets represent a powerful framework for complete estimation of type-I and type-II errors in Hi-C benchmarks (Jorge et al., 2025).

Two perspectives deserve emphasis. First, while HiCDOC has so far been applied to the classical two-compartment (A/B) framework, its clustering-based method is in principle compatible with detecting more than two chromatin states. This feature provides an original approach to explore finer levels of chromatin modification that cannot be captured using PCA-based methods, like subcompartment dynamics for instance. Second, although our evaluations and our tests focused on pairwise comparisons between two experimental groups, HiCDOC is inherently compatible with more complex experimental designs.

In summary, HiCDOC introduces a practical and robust method for differential analysis of the chromatin compartment. By leveraging replicate information, providing extensive quality-control metrics, and achieving strong concordance with both gene expression and ground truth data, HiCDOC expands the computational toolkit available for studying chromatin structure. We anticipate that HiCDOC will help to build a more comprehensive picture of 3D genome organization and its links to cellular function.

## Acknowledgements

The authors thank the Chrocogen community and the Genotoul-Bioinfo platform (https://doi.org/10.15454/1.5572369328961167E12) for providing valuable assistance and resources. Founding was partly provided by the DIGIT-BIO INRAE Metaprogram “ChrocoNET”.

## Author contributions

C.K., E.M., and M.Z. wrote the package. S.F., E.M., and M.Z. wrote the paper. M.Z. and S.F. supervised the study. All authors designed the method and analyzed the data. All authors have read and approved the article.

## Data, script, code, and supplementary information availability

Benchmarking scripts and additional results are available online at https://elisemaigne.pages-forge.inrae.fr/benchmark_hicdoc/.

